# *Drosophila Nepl15* controls glycogen and lipid storage by modulating insulin signaling

**DOI:** 10.64898/2026.07.15.738827

**Authors:** Shahira Helal Arzoo, Surya Jyoti Banerjee

## Abstract

Deletion of the *Drosophila*-specific gene *Neprilysin-like 15* (*Nepl15*) results in significant reductions in glycogen and glycerolipid storage in mutant males and increased glycogen storage in mutant females without affecting food intake. Previous studies also indicated downregulation of insulin/mTOR signaling, the central pathway regulating nutrient homeostasis, growth, fertility, locomotor activity, and lifespan. Consistent with these metabolic alterations, *Nepl15* mutants exhibit several anti-obesity and healthy-aging phenotypes but display markedly reduced survival under starvation, suggesting impaired nutrient reserve utilization. The present study investigated the intracellular mechanisms underlying these phenotypes by examining insulin signaling and the expression of genes involved in carbohydrate and lipid metabolism. Although transcript levels of the insulin-like peptides (*Dilp2*, *Dilp5*, and *Dilp6*) and the insulin receptor (*InR*) remained unchanged, *Nepl15* mutants exhibited reduced Akt phosphorylation, increased dFoxo abundance, reduced *Glut1* expression, and previously reported suppression of mTOR signaling, indicating attenuation of insulin signaling downstream of the insulin receptor. Consistent with impaired anabolic signaling, male mutants showed reduced expression of glycogen metabolic genes (*GlyS* and *GlyP*), whereas genes involved in lipid metabolism exhibited predominantly male-specific reductions, including *Lipin*, *Acc*, *Fasn1*, and *Fasn2*, while *Midway* and *Brummer* remained unchanged. In addition, expression of the metabolic regulator *PGC1*α (*spargel*) was significantly reduced in males, consistent with previously reported reductions in *AMPK*α expression. Together, these findings demonstrate that *Nepl15* functions as an upstream regulator of insulin-dependent anabolic signaling and nutrient partitioning, linking intracellular signaling to glycogen and lipid storage independently of nutrient intake. These results identify *Nepl15* as a previously unrecognized regulator of metabolic homeostasis in *Drosophila* and provide new insight into the metabolic functions of the evolutionarily conserved neprilysin family.

## Introduction

Neprilysins (*Nep*s) are zinc (II)-dependent M13 family metalloendopeptidases, structurally and functionally related to endothelin-converting enzymes, PHEX, KELL, and other family members, and conserved from mammals to invertebrates (Bland, Pinney, Thomas, Turner, & Isaac, 2008; Meyer, Panz, Zmojdzian, Jagla, & Paululat, 2009; Nalivaeva & Turner, 2013; Sitnik et al., 2014; Turner, Isaac, & Coates, 2001). Typical mammalian *Nep*s are type II integral membrane glycoproteins bearing conserved catalytic motifs (HEXXH, EXXAD) and substrate-binding and maturation motifs (NAY/FY, CXXW), and they require Zn(II) coordination by conserved His and Glu residues to hydrolyze small peptide substrates, generally cleaving N-terminal to hydrophobic residues (Devault et al., 1988; Nalivaeva & Turner, 2013; Oefner, D’Arcy, Hennig, Winkler, & Dale, 2000; Turner, Brown, Carson, & Barnes, 2000). *Nep*s are broadly expressed across kidney, brain, lung, reproductive tissue, adipocytes, and smooth muscle, where they degrade peptides including enkephalins, natriuretic peptides, bradykinin, angiotensins, and, critically for metabolism, insulin B-chain, glucagon, and glucagon-like peptide-1 (Nalivaeva & Turner, 2013; Roques, Noble, Daugé, Fournié-Zaluski, & Beaumont, 1993; Trebbien et al., 2004; Willard, Barrow, & Zraika, 2017).

This capacity to degrade metabolic hormones directly implicates *Nep*s in insulin signaling and glucose homeostasis. Elevated plasma *Nep* levels are observed in obese and overweight individuals, and *Nep* inhibition in obese rats increases circulating glucose and tissue glucose uptake, indicating that elevated *Nep* activity exacerbates insulin resistance (Standeven et al., 2011). Hyperglycemic and high-fat conditions themselves induce *Nep* activity: excess saturated fatty acids and glucose elevate *Nep* activity in cultured endothelial cells and mouse islets, coinciding with oxidative stress and impaired insulin secretion, while diet-induced obesity increases *Nep* levels in both plasma and tissue (Muangman, Spenny, Tamura, & Gibran, 2003; Standeven et al., 2011; Zraika et al., 2013). Yet the relationship is not strictly linear. *Nep*-deficient mice on a high-fat diet show improved glucose clearance and elevated GLP-1 despite greater weight gain, and heterozygous *Nep* mutants also show age-related weight gain (Becker et al., 2010; Willard et al., 2017), suggesting *Nep* activity and nutrient status are coupled through a more complex regulatory loop than simple induction or suppression.

In *Drosophila*, 24 *Nep*rilysin-like genes are distributed across five phylogenetic clades, reflecting partial conservation with and partial divergence from mammalian *Nep*s (Bland et al., 2008; Turner et al., 2001). Several fly *Nep*s have documented roles relevant to metabolism and reproduction: *Nep*2 cleaves amyloid-beta and tachykinin (Finelli, Kelkar, Song, Yang, & Konsolaki, 2004; Thomas et al., 2005), Nep4 cleaves substance P and angiotensin I and regulates dilp expression, food intake, body size, and viability when overexpressed (Hallier et al., 2016; Meyer et al., 2009), Goe regulates spermiogenesis (Matsuoka, Gupta, Suzuki, Hiromi, & Asaoka, 2014), and Frma regulates female receptivity (Findlay et al., 2014). *Nep*1 expression is upregulated by a high-fat diet (Heinrichsen et al., 2014), and genome-wide knockdown screening identified variable effects of individual *Nep*s on triglyceride content (Pospisilik et al., 2010). Notably, *Nepl15* showed no transcriptional change under high-fat diet and was not captured in prior obesity or glucome screens (Heinrichsen et al., 2014; Pospisilik et al., 2010; Ugrankar et al., 2015), leaving its metabolic function entirely undefined despite the broader family’s clear links to insulin signaling and nutrient disorders.

*Drosophila* is a powerful model for metabolic disease because its neuroendocrine control of nutrient homeostasis centered on *Drosophila* insulin-like peptides (Dilps) and adipokinetic hormone (Akh), the functional analog of glucagon, is mechanistically conserved with the mammalian insulin/glucagon and TOR axes (Baker & Thummel, 2007; Kannan & Fridell, 2013; Padmanabha & Baker, 2014). Dietary sugars and lipids, absorbed by midgut enterocytes, enter the hemolymph and are taken up by the fat body, the fly’s functional liver/adipose equivalent, where glucose is converted via UDP-glucose into trehalose and glycogen, and absorbed diacylglycerol is esterified into triglyceride and stored in lipid droplets (Arrese & Soulages, 2010; Kühnlein, 2012; Matsuda, Yamada, Yoshida, & Nishimura, 2015; Padmanabha & Baker, 2014).

Insulin signaling in flies begins with Dilps 2, 3, and 5, secreted from brain insulin-producing cells (IPCs), activating the single insulin receptor (InR) and its substrate Chico, which recruits PI3K, generates PIP3, and activates Akt via PDK1 (Baker & Thummel, 2007; Fernandez, Tabarini, Azpiazu, Frasch, & Schlessinger, 1995; Kim & Rulifson, 2004; Nässel, 2012; Rulifson, Kim, & Nusse, 2002). Active Akt phosphorylates dFoxo, excluding it from the nucleus and blocking transcription of growth-inhibitory genes including 4E-BP, while simultaneously promoting glucose transporter recruitment, glucose uptake, and glycogen synthesis (Teleman, Chen, & Cohen, 2005a, 2005b). Akt also activates Rheb and inhibits the TSC1/TSC2 complex to stimulate TOR, which drives ribosomal biogenesis and translation via S6 kinase and represses 4E-BP, together promoting nutrient absorption, glycogen and lipid storage, and growth while suppressing glycolysis and lipolysis (Danielsen, Moeller, & Rewitz, 2013; Grönke, Clarke, Broughton, Andrews, & Partridge, 2010; Mirth & Riddiford, 2007; Teleman et al., 2005a). Akh, secreted in response to low nutrient states, opposes this anabolic program by mobilizing stored glycogen and lipid, mirroring glucagon’s counter-regulatory role in mammals (Van der Horst, 2003).

These pathways are highly sensitive to genetic and dietary perturbation, producing phenotypes that model human metabolic disease. Ablation of IPCs or triple-null mutation of dilps 2, 3, and 5 elevates circulating glucose, trehalose, glycogen, and lipid (Broughton et al., 2005; Grönke et al., 2005; Rulifson et al., 2002), and individual dilp knockdowns similarly raise triglyceride levels, indicating additional insulin-independent inputs into storage regulation (Pospisilik et al., 2010). High-fat and high-sugar diets (HFD/HSD) likewise disrupt this homeostasis, altering *Nep* and insulin pathway activity and nutrient storage in ways that recapitulate mammalian obesity and diabetes phenotypes (Heinrichsen et al., 2014; Musselman & Kühnlein, 2018). This conservation from receptor to downstream Akt/Foxo/TOR effectors makes *Drosophila* genetic and dietary models directly relevant to dissecting how individual gene products, such as *Nepl15*, tune nutrient partitioning through these same conserved nodes.

Deletion of *Nepl15*, a *Drosophila*-specific gene with no prior functional characterization in metabolism, produces a pronounced and sexually dimorphic storage phenotype: mutant males show significant reductions in glycogen and glycerolipid stores, whereas mutant females show a significant increase in glycogen storage, with neither sex showing any change in food intake relative to controls. This dissociation between feeding behavior and nutrient storage indicates that *Nepl15* acts on the intracellular routing or partitioning of absorbed nutrients rather than on appetite or ingestion (Banerjee et al., 2021). Recent studies further point to downregulation of insulin/mTOR signaling, the central pathway integrating nutrient and energy homeostasis with growth, fertility, motor activity, and lifespan, in *Nepl15* mutants of both sexes (Arzoo et. Al, 2026). Consistent with reduced anabolic signaling, mutant flies display multiple anti-obese and anti-aging physiological and cellular traits, yet they also show severe mortality specifically under starvation (Banerjee et al., 2021), suggesting that the same signaling changes that confer lean, youthful phenotypes under fed conditions come at the cost of nutrient reserves needed to survive fasting.

The present study investigates the intracellular signaling pathways underlying these phenotypes in both male and female *Nepl15* mutant flies to determine whether their altered nutrient storage results from impaired anabolic signaling rather than changes in nutrient intake. We examined key regulators of insulin signaling together with genes involved in glycogen metabolism, lipid metabolism, and cellular energy homeostasis. Although transcript levels of the insulin-like peptides (*Dilp2*, *Dilp5*, and *Dilp6*) and the insulin receptor (*InR*) remained unchanged in both sexes, *Nepl15* mutants exhibited reduced Akt phosphorylation, increased dFoxo abundance, reduced *Glut1* expression in males, and previously reported suppression of mTOR signaling (Arzoo et al., 2026), indicating attenuation of insulin signaling downstream of the insulin receptor. Consistent with these signaling changes, male mutants displayed reduced expression of genes involved in glycogen metabolism (*GlyS* and *GlyP*) and de novo lipogenesis (*Acc*, *Fasn1*, and *Fasn2*), whereas females exhibited more limited transcriptional changes, including reduced *Lipin* and *Fasn1* expression and increased *Acc* expression. Expression of *Midway* and *Brummer* remained unchanged in both sexes. In addition, the metabolic regulator *PGC1*α (*spargel*) was significantly reduced in males, consistent with our previous observation of decreased *AMPK*α expression (Arzoo et al., 2026). Together, these findings identify *Nepl15* as an upstream regulator of insulin-dependent anabolic signaling and nutrient partitioning that links intracellular signaling to sex-specific regulation of glycogen and lipid metabolism despite normal nutrient intake.

## Methods

### Fly Husbandry and Experimental Cohorts

Isogenic *Nepl15* knockout flies (*w^1118^; Nepl15^KO^*, hereafter referred to as *Nepl15^KO^*) generated by ends-out homologous recombination as previously described (Banerjee et al., 2021)were used throughout this study. The isogenic w^1118^ line served as the genetic background control.

All experiments were performed using age-matched male flies at 5–7 days. To ensure consistency across genotypes, flies were maintained on identical food under the same population density and cohort conditions. Parental crosses were established by placing 10 virgin females and 5 males per vial (Genesee Scientific 32-113RL). Vials were maintained in a Percival Scientific incubator (DR-36NL) at 25 °C and 70% relative humidity on Nutri-Fly Pre-Mixed Fly Food (Genesee Scientific 66-113) supplemented with 20mL 20% Tegosept (methyl 4-hydroxybenzoate, Sigma H3647) and 20mL propionic acid (Ward’s Science 470302-290) per 3.5L of food to minimize fungal and bacterial contamination. To collect virgin progeny, parental vials were maintained at 18 °C, and newly eclosed flies were collected within 16 hours of eclosion. Progeny were sexed under light CO□ anesthesia and transferred to fresh food for aging at approximately 20 flies per vial. Flies were transferred to fresh food two to three times per week to maintain consistent nutritional conditions. All experiments were conducted under controlled environmental conditions to minimize variability due to diet, density, or age.

### Western Blot

Whole-body protein lysates were prepared from adult male flies (three independent biological replicates; 20 flies per replicate). Flies were homogenized on ice in RIPA lysis buffer (Sigma 20-188) supplemented with Halt protease and phosphatase inhibitor (Thermo Scientific1861280) and β-mercaptoethanol (Sigma M3148), denatured at 95 °C for 5 min, and centrifugation at 4 °C. Protein concentration was determined by Bradford assay (Bio-Rad 5000205) following 1:4 sample dilution, and equal amounts of protein (30 µg per lane) were resolved by SDS–PAGE (Bio-Rad 4568096) and transferred onto methanol-activated PVDF membranes (Bio-Rad 1620177) using Tris–glycine transfer buffer (25 mM Tris, 192 mM glycine, 20% methanol). Membranes were blocked in TBS containing 5% milk (or BSA for phospho-proteins) and incubated overnight at 4°C with primary antibodies against Rabbit polyclonal anti-*Drosophila* phospho-AKT (Cell Signaling Technology 4054), Rabbit polyclonal anti-total AKT (Cell Signaling Technology 9272) (van Dam et al., 2020), rabbit anti-pFoxo1 (Cell Signaling Technology 9461,), rabbit anti-dFoxo1 (Abcam ab195977) (Pandey et al., 2023). After washing in TBST, membranes were incubated with HRP-conjugated secondary antibodies (Jackson ImmunoResearch anti-rabbit IgG 111-035-047) and developed using chemiluminescence (Western ECL Substrate, BioRad 1705062). All targets were normalized to loading control (primary Mouse anti-tubulin, DHSB E7). Relative protein levels were compared relative to control samples.

### Quantitative RT–PCR

Quantitative PCR (qPCR) primers were either designed using the Integrated DNA Technologies PrimerQuest Tool (Kalendar, 2025) or adopted from previously published studies. All primers (Table 1) were validated for target specificity and efficiency before use. Total RNA was isolated from the indicated tissues using TRI Reagent (Sigma-Aldrich T9424), followed by cDNA synthesis with iScript Reverse Transcription Supermix (Bio-Rad 1708841). Gene expression analysis was performed using SsoAdvanced Universal SYBR Green Supermix (Bio-Rad 1725271) on a CFX Duet Real-Time PCR Detection System (Bio-Rad). Transcript abundance of target genes was normalized to the reference gene *RpL32* (Banerjee et al., 2021; Joshi, Banerjee, Curtiss, & Ashley, 2022), and relative expression levels were calculated using the 2^−ΔΔCt^ method. All procedures and data analyses were conducted in accordance with the Minimum Information for Publication of Quantitative Real-Time PCR Experiments (MIQE) guidelines (Banerjee et al., 2021).

**Table 1:**
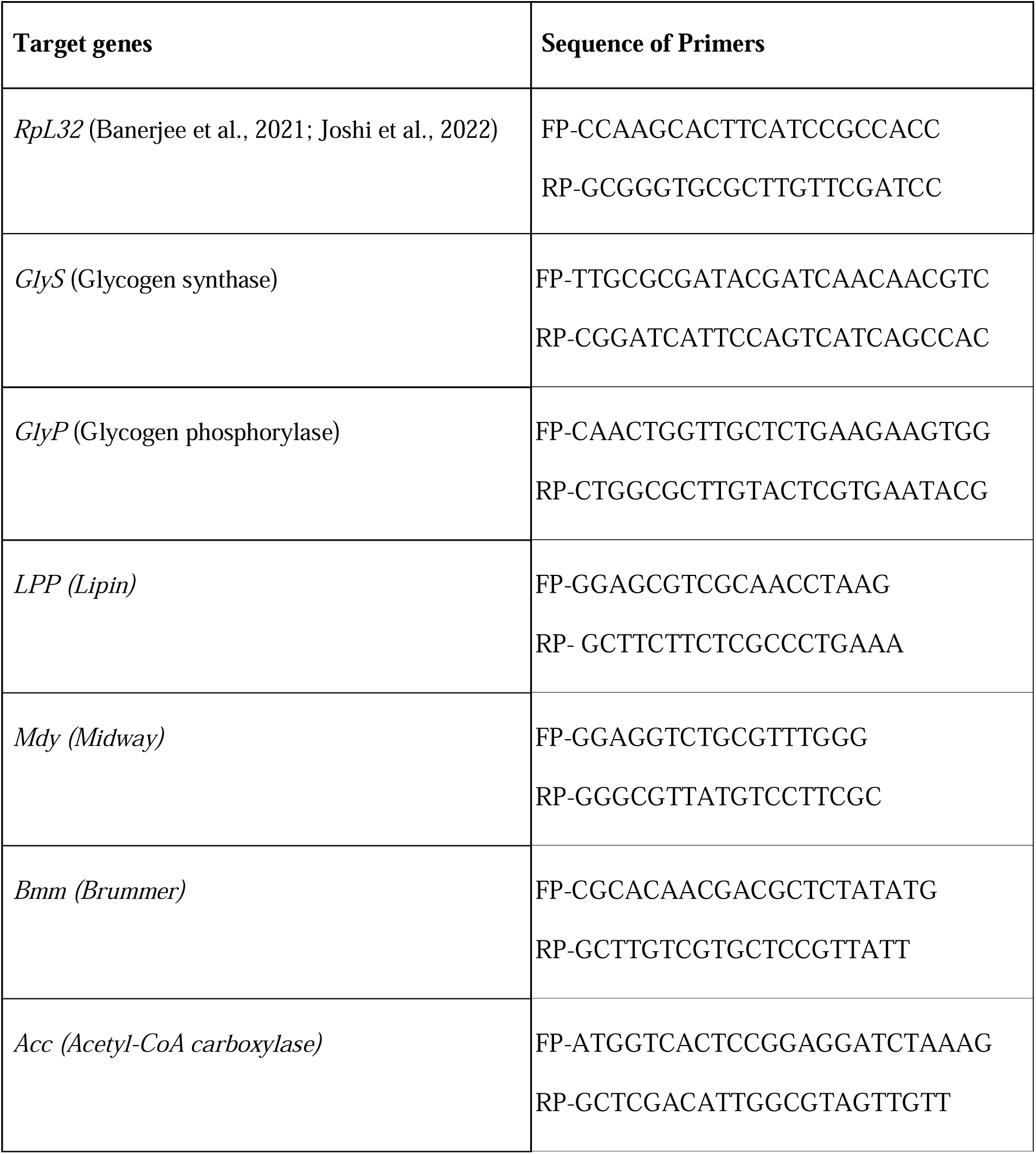
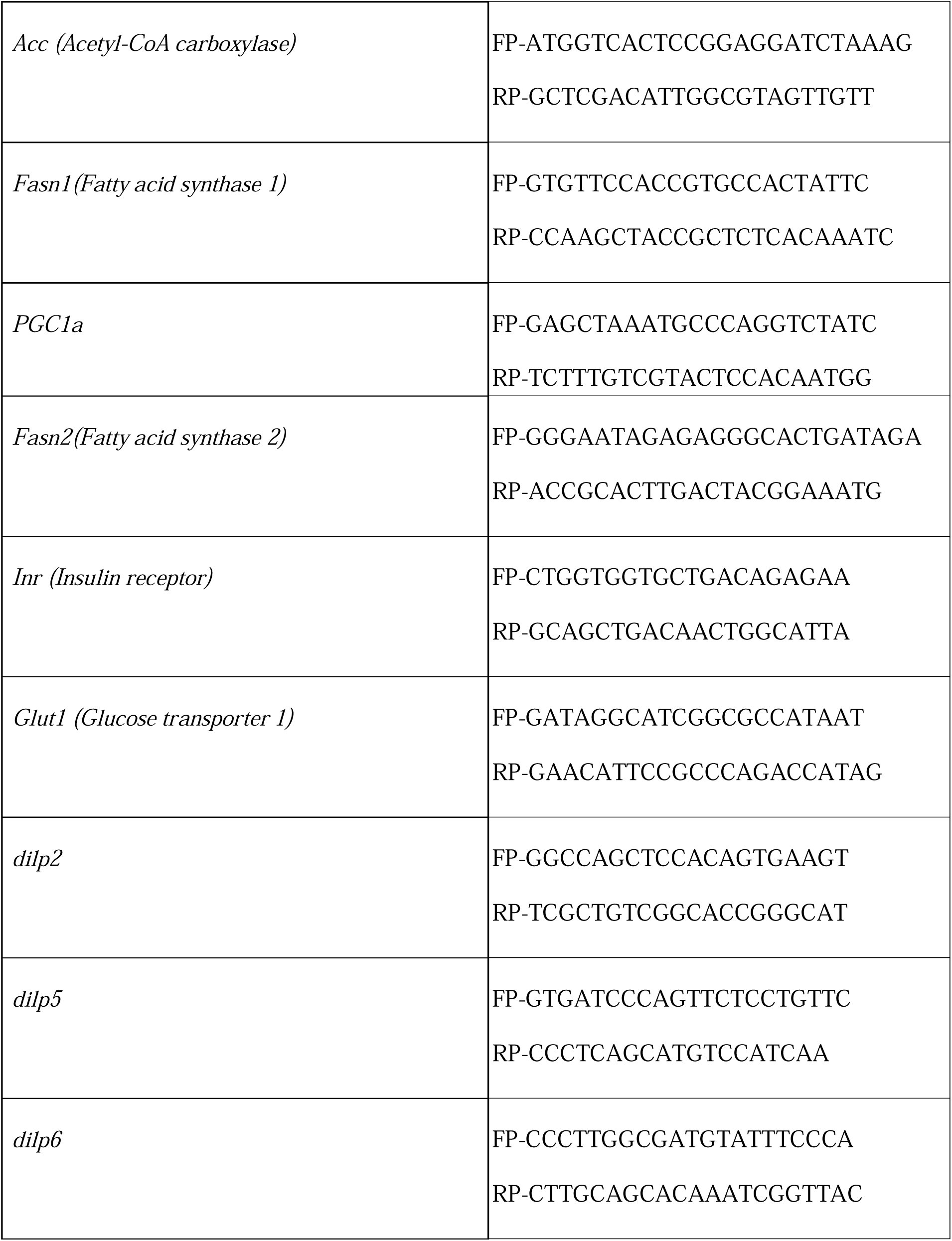
Primer sequences used for qRT-PCR analysis. Forward (FP) and reverse (RP) primer sequences are listed:

### Statistical analysis

All statistical analyses were performed using GraphPad Prism (version 9). Data are presented as mean ± standard error of the mean (SEM) from three independent biological replicates. Comparisons between control and *Nepl15* knockout flies within each sex were performed using an unpaired two-tailed Student’s *t*-test. Differences were considered statistically significant at P < 0.05. Statistical significance is indicated as *P* < 0.05 (*), P < 0.01 (), P < 0.001 (*), and *P* < 0.0001 (****), whereas ns denotes not significant (*P* ≥ 0.05).

## Result

### 1. *Nepl15* loss attenuates insulin signalling in flies

**Figure 1:**
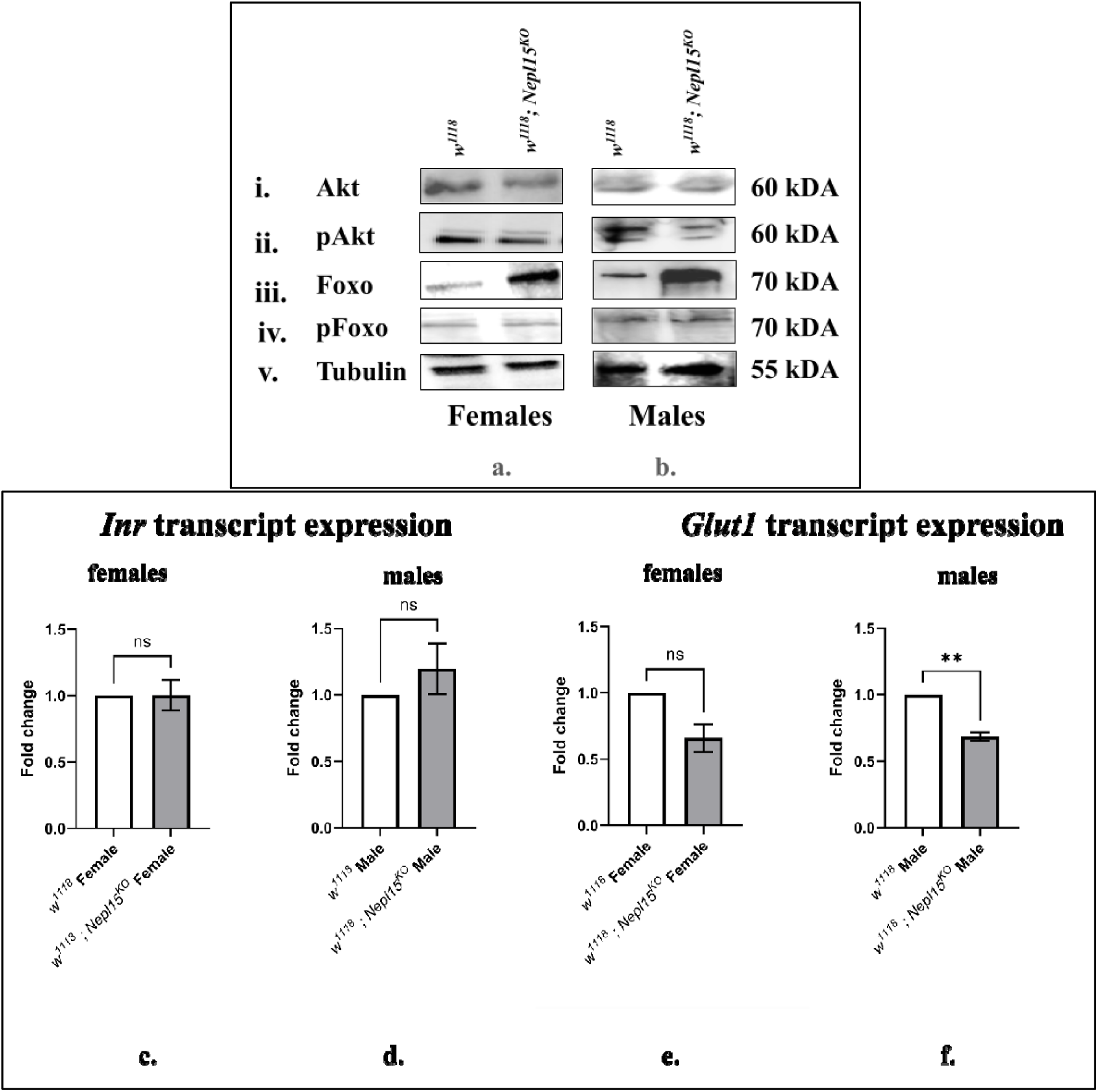
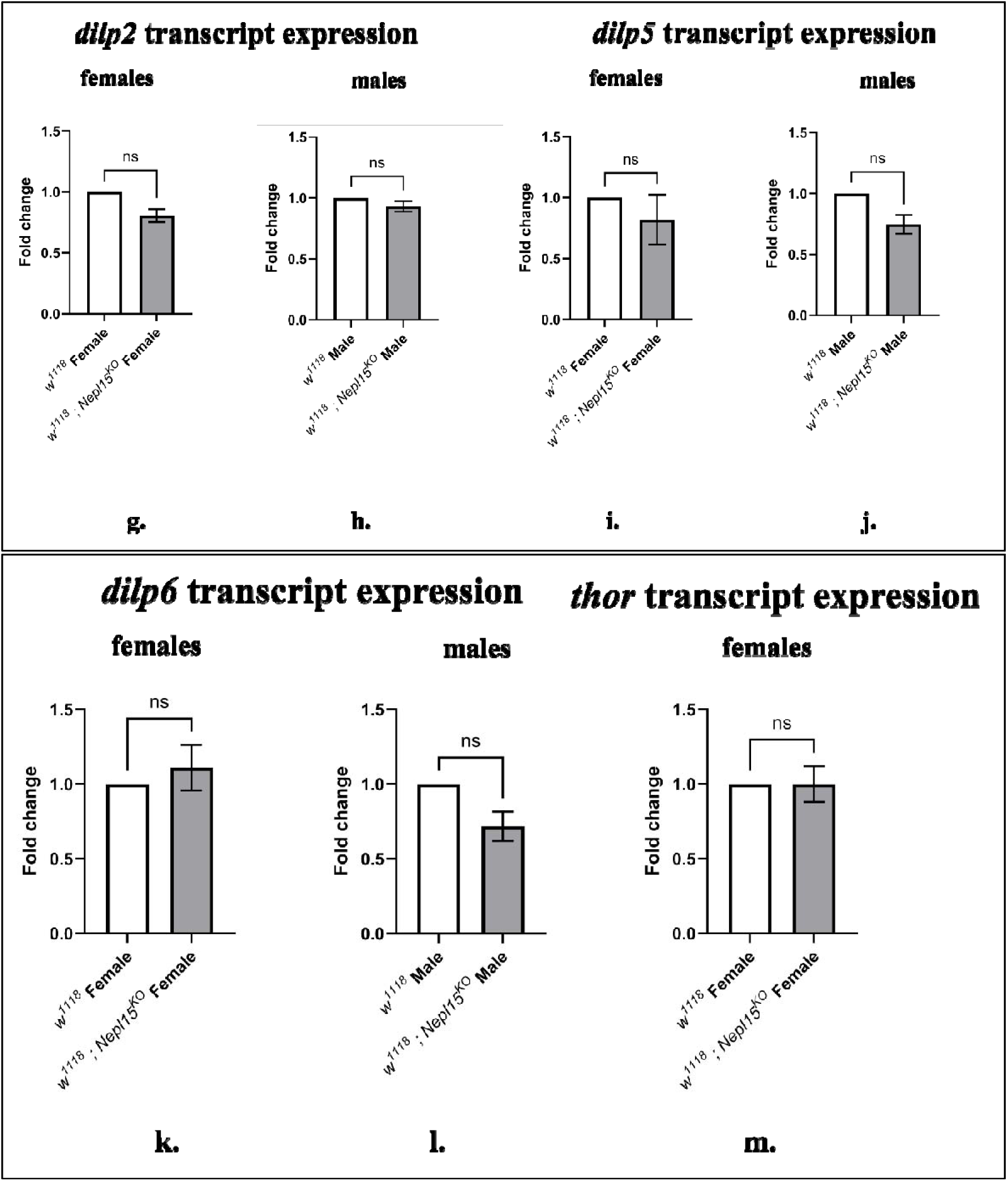
Loss of *Nepl15* alters components of the insulin signaling pathway in adult *Drosophila*. (a, b) Representative Western blot analysis of whole-body protein lysates from 5–7-day-old female (a) and male (b) w^1118^ control and *Nepl15*^KO^ flies. Membranes were probed with antibodies against i. total Akt (∼60 kDa), ii. phospho-Akt (pAkt; Ser505, ∼60 kDa), iii. total dFoxo (∼70 kDa), iv. phospho-dFoxo (pFoxo; ∼70 kDa), and v. tubulin (∼55 kDa), which served as the loading control. Molecular weights (kDa) are indicated on the right. (c–f) Relative transcript levels of the insulin receptor (InR) (c, d) and glucose transporter 1 (*Glut1*) (e, f) in female (c, e) and male (d, f) flies determined by quantitative RT-qPCR. InR transcript levels were not significantly different between genotypes in either sex, whereas *Glut1* expression was significantly reduced in *Nepl15* knockout males but not females. (g–l) Relative transcript levels of *Dilp2* (g, h), *Dilp5* (i, j), and *Dilp6* (k, l) in female (g, i, k) and male (h, j, l) flies. Expression of all three insulin-like peptide genes was not significantly altered in *Nepl15* knockout flies. (m) Relative transcript abundance of *Thor* in female flies showing no significant difference between genotypes. Bars represent mean ± SEM from three independent biological replicates. Statistical significance was determined using an unpaired two-tailed Student’s t-test. ns, not significant; **, P < 0.01.

Because insulin signaling is the principal regulator of glycogen and lipid synthesis in *Drosophila* melanogaster (Baker & Thummel, 2007; Teleman et al., 2005a), we investigated whether the reduced nutrient storage previously observed in *Nepl15* knockout flies (Banerjee et al., 2021) was associated with alterations in the insulin signaling pathway. We first examined transcript levels of the insulin-like peptides *Dilp2*, *Dilp5*, and *Dilp6*, which are major regulators of systemic insulin signaling in adult flies. No significant differences in the expression of *Dilp2*, *Dilp5*, or *Dilp6* were detected between *Nepl15* knockout and control flies in either females or males (Fig. 1 g – l), suggesting that loss of *Nepl15* does not affect insulin-like peptide expression.

To determine whether insulin signaling was altered at the receptor level, we measured transcript abundance of the insulin receptor (InR). InR expression was comparable between control and *Nepl15* knockout flies in both sexes (Fig. 1 c, d), indicating that the reduced nutrient storage phenotype is unlikely to result from altered insulin receptor expression. In contrast, expression of the glucose transporter *Glut1*, a downstream target that facilitates insulin-dependent glucose uptake, was significantly reduced in *Nepl15* knockout males but remained unchanged in females (Fig. 1 e, f). Reduced *Glut1* expression in males is consistent with decreased cellular glucose uptake and provides a potential mechanism contributing to the reduced glycogen accumulation previously reported in *Nepl15* mutant flies (Banerjee et al., 2021).

To further investigate intracellular insulin signaling, we analyzed key components of the Akt–dFoxo signaling axis by Western blot (Fig. 1 a, b). Total Akt protein abundance was similar between controls and *Nepl15*^KO^ in both sexes. However, phosphorylated Akt (pAkt), the active form of Akt, was reduced in *Nepl15*^KO^ flies, with the reduction being more pronounced in males. Conversely, total dFoxo protein abundance was increased, whereas phosphorylated dFoxo (pFoxo) levels were reduced, particularly in male mutants. Because Akt-mediated phosphorylation inhibits dFoxo activity (Giannakou et al., 2004; Teleman et al., 2005a), these observations are consistent with reduced insulin pathway activation following loss of *Nepl15*. These findings further extend our previous study demonstrating significantly reduced mTOR expression (Arzoo et. al, 2026) together with increased *Thor* expression in *Nepl15*^KO^ flies (Banerjee et al., 2021). *Thor* transcript levels were not significantly altered in female mutants in the present study (Fig. 1 m), indicating that downstream activation of this translational regulator is more pronounced in males. Collectively, the unchanged expression of *Dilp2*, *Dilp5*, *Dilp6*, and InR, together with reduced *Glut1* expression, decreased Akt phosphorylation, increased dFoxo abundance, and our previously reported reduction in mTOR signaling, indicate that loss of *Nepl15* suppresses insulin-dependent anabolic signaling primarily downstream of the insulin receptor. This reduction in anabolic signaling provides a molecular framework for the impaired glycogen and lipid storage observed in *Nepl15* mutant flies (Banerjee et al., 2021).

### 2. *Nepl15* loss suppresses glycogen metabolism in male flies

**Figure 2.**
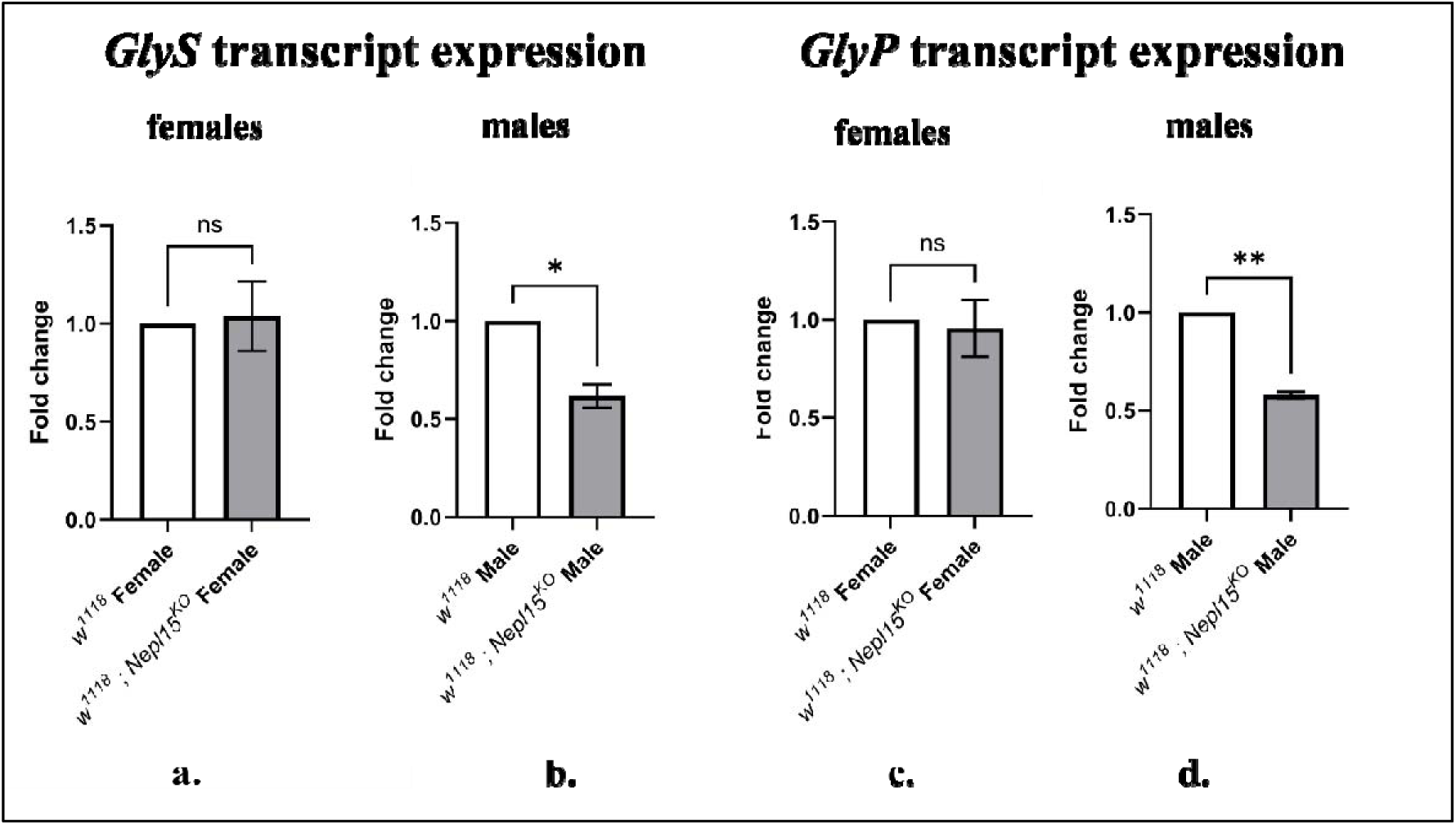
Loss of *Nepl15* reduces the expression of glycogen metabolic genes in male *Drosophila*. Relative transcript levels of *GlyS* (glycogen synthase) (a, b) and *GlyP* (glycogen phosphorylase) (c, d) were determined by quantitative RT-PCR in female (a, c) and male (b, d) control and *Nepl15*^KO^ flies. *GlyS* expression was significantly reduced in male *Nepl15*^KO^ flies but was unchanged in females. Similarly, *GlyP* transcript levels were significantly reduced in male *Nepl15*^KO^ flies, whereas no significant difference was observed in females. Bars represent mean ± SEM from three independent biological replicates. Statistical significance was determined using an unpaired two-tailed Student’s t-test. P < 0.05 (*), P < 0.01 (**), ns, not significant.

Because insulin signaling promotes glycogen storage by regulating both glycogen synthesis and glycogen mobilization (Baker & Thummel, 2007; Teleman et al., 2005b), we next examined whether loss of *Nepl15* alters the expression of genes involved in glycogen metabolism. Quantitative RT-PCR was performed to determine the transcript abundance of *GlyS*, which encodes glycogen synthase, the rate-limiting enzyme responsible for glycogen synthesis, and *GlyP*, which encodes glycogen phosphorylase, the enzyme that catalyzes glycogen degradation. Expression of *GlyS* was not significantly altered in female *Nepl15* knockout flies compared with controls (Fig. 2 a). In contrast, male mutants exhibited a significant reduction in *GlyS* transcript abundance (Fig. 2 b). Similarly, *GlyP* expression remained unchanged in females (Fig. 2 c) but was significantly reduced in male *Nepl15* knockout flies relative to controls (Fig. 2 d). The coordinated reduction of both *GlyS* and *GlyP* expression in males indicates that loss of *Nepl15* alters transcriptional regulation of glycogen metabolism in a sex-specific manner. These findings are consistent with the reduced insulin-dependent anabolic signaling observed in *Nepl15* mutants (Fig. 1 b) and with our previous report demonstrating decreased whole-body glycogen storage in male *Nepl15* knockout flies (Banerjee et al., 2021). Together, these data suggest that suppression of insulin signaling in the absence of *Nepl15* is accompanied by reduced expression of key enzymes governing glycogen homeostasis.

### 3. *Nepl15* loss alters the expression of genes involved in lipid metabolism

**Figure 3.**
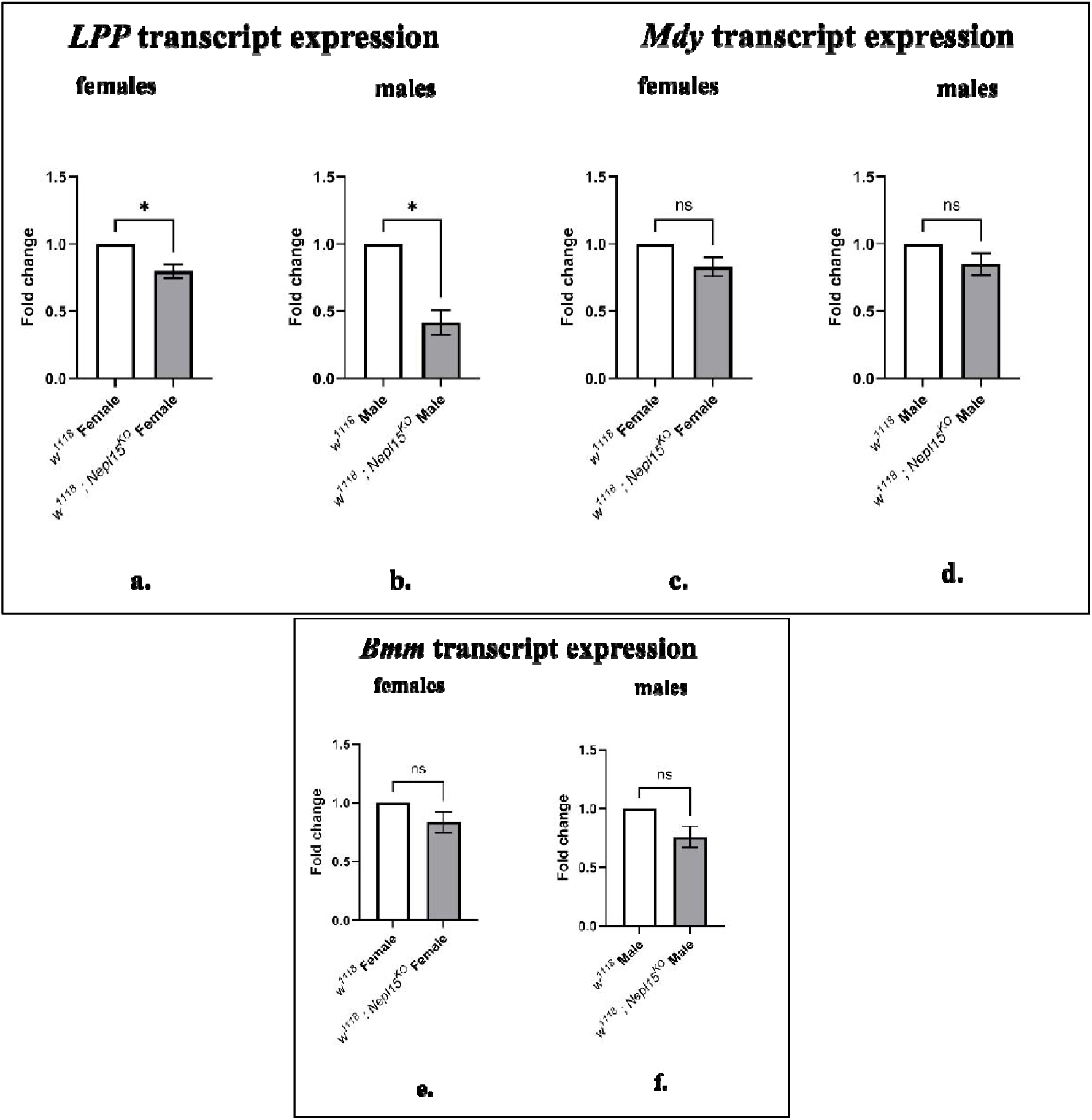
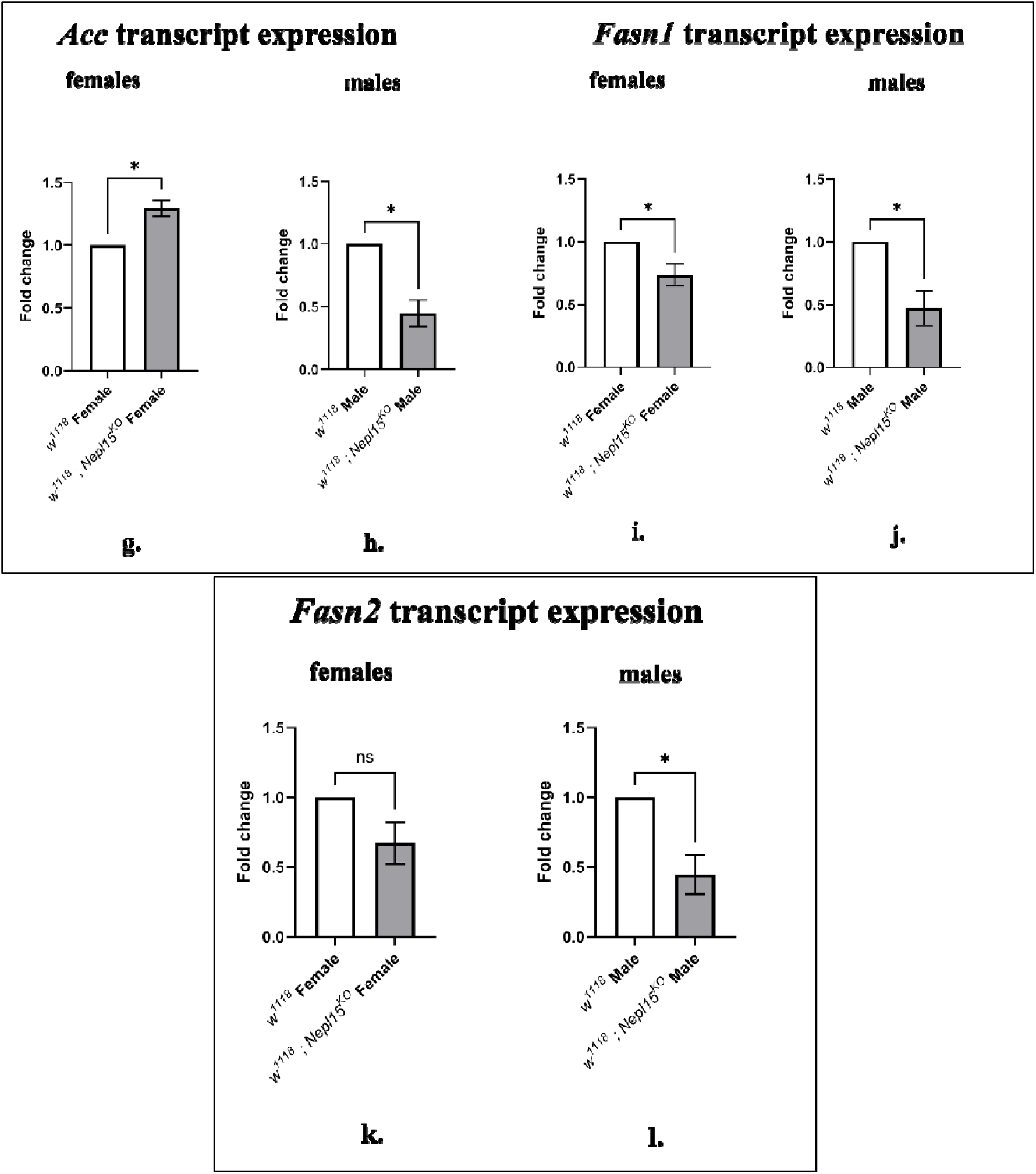
Loss of *Nepl15* alters the expression of lipid metabolism genes in adult *Drosophila*. Relative transcript levels of *Lpp (Lipin)* (a, b), *Mdy (Midway)* (c, d), *Bmm (Brummer)* (e, f), *Acc (acetyl-CoA carboxylase)* (g, h), *Fasn1 (fatty acid synthase 1)* (i, j), and *Fasn2 (fatty acid synthase 2)* (k, l) were determined by quantitative RT-PCR in female (a, c, e, g, i, k) and male (b, d, f, h, j, l) control and *Nepl15^KO^* flies. *Lpp* expression was significantly reduced in both female and male *Nepl15^KO^* flies. *<u>Mdy</u>* and *Bmm* transcript levels were not significantly altered in either sex. *Acc* expression was significantly increased in female mutants but significantly reduced in male mutants. *Fasn1* transcript abundance was significantly reduced in both female and male *Nepl15*^KO^ flies, whereas *Fasn2* expression was significantly reduced only in males and was unchanged in females. Bars represent mean ± SEM from three independent biological replicates. Statistical significance was determined using an unpaired two-tailed Student’s t-test. *P* < 0.05 (*), *P* < 0.01 (**), ns, not significant

Because insulin–mTOR signaling regulates lipid synthesis, storage, and mobilization in *Drosophila* (Baker & Thummel, 2007; Kühnlein, 2012; Teleman et al., 2005a), we next investigated whether the suppression of insulin-dependent anabolic signaling in *Nepl15* knockout flies was accompanied by altered expression of genes involved in lipid metabolism. We examined transcripts encoding enzymes involved in triglyceride synthesis (*Mdy*), fatty acid synthesis (*Acc*, *Fasn1*, and *Fasn2*), lipid mobilization (*Bmm*), and the phosphatidic acid phosphatase Lipin (*Lipin*), a central regulator of glycerolipid metabolism.

Quantitative RT-PCR analysis revealed a significant reduction in *Lipin* transcript abundance in both female and male *Nepl15* knockout flies relative to controls (Fig. 3 a, b). In contrast, expression of *Mdy* and *Bmm* was not significantly altered in either sex (Fig. 3 c – f), indicating that neither diacylglycerol acyltransferase-mediated triglyceride synthesis nor Brummer-dependent lipolysis was transcriptionally affected by loss of *Nepl15*. The expression of genes involved in de novo fatty acid synthesis exhibited marked sex-specific differences. *Acc* transcript abundance was significantly increased in female *Nepl15* knockout flies but significantly reduced in males (Fig. 3 g, h). Similarly, *Fasn1* expression was significantly reduced in both sexes, although the reduction was more pronounced in males (Fig. 3 i, j). *Fasn2* transcript levels were significantly reduced in male mutants but remained unchanged in females (Fig. 3 k, l). Collectively, these findings demonstrate that loss of *Nepl15* selectively alters the transcription of genes involved in lipid metabolism, with the strongest effects observed in males. The coordinated reduction of *Lipin*, *Acc*, *Fasn1*, and *Fasn2* expression in male *Nepl15* knockout flies is consistent with reduced lipogenic capacity and agrees with the suppression of insulin-dependent anabolic signaling described above (Fig. 1). These transcriptional changes further support our previous observation that *Nepl15* deficiency results in reduced triglyceride storage (Banerjee et al., 2021).

### 4. *Nepl15* loss reduces expression of metabolic regulators in male flies

**Figure 4.**
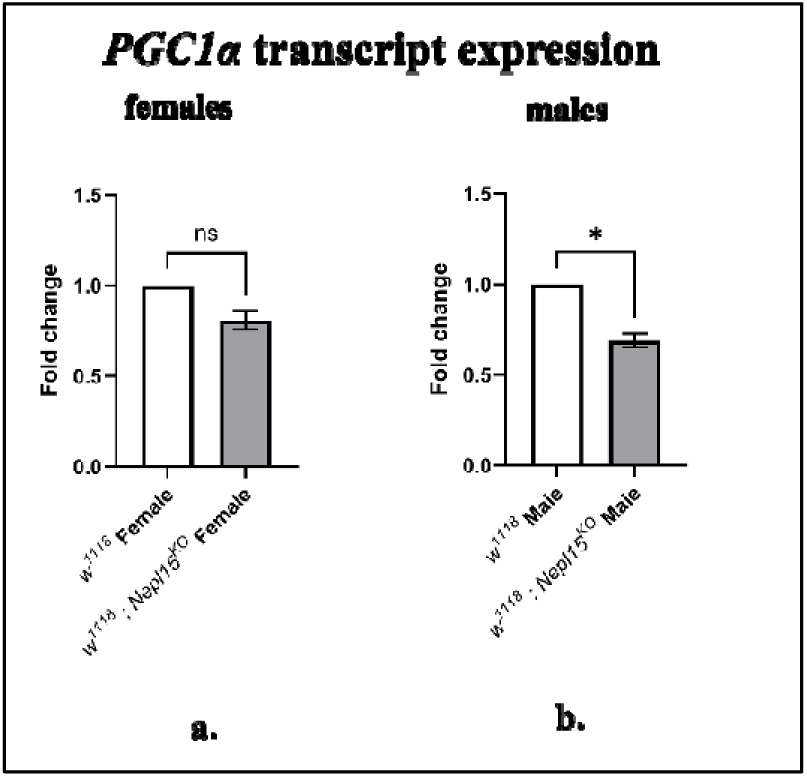
Loss of *Nepl15* reduces *PGC1*_α_ (*Spargel*) transcript expression in male *Drosophila*. Relative transcript levels of *PGC1*_α_ (a, b) were determined by quantitative RT-PCR in female (a) and male (b) control and *Nepl15^KO^* flies. *PGC1*_α_ expression was significantly reduced in male *Nepl15^KO^*flies but was unchanged in females. Bars represent mean ± SEM from three independent biological replicates. *P* < 0.05 (*), ns, not significant.

Because insulin and AMP-activated protein kinase (AMPK) signaling coordinately regulate cellular energy metabolism through the transcriptional coactivator PGC1α, we next examined whether loss of *Nepl15* altered *PGC1*_α_ expression. Quantitative RT-PCR analysis revealed that *PGC1*_α_ transcript abundance was not significantly different between female *Nepl15^KO^* flies and controls (Fig. 4 a). In contrast, male *Nepl15^KO^* flies exhibited a significant reduction in *PGC1*_α_ expression relative to controls (Fig. 4 b). These findings are consistent with our previous study demonstrating significantly reduced *AMPK*_α_ expression in both female and male *Nepl15* knockout flies (Arzoo et. al, 2026). Together, the reduction in *AMPK*_α_ and the male-specific decrease in *PGC1*_α_ expression suggest that loss of *Nepl15* is associated with altered transcriptional regulation of cellular energy metabolism in addition to the suppression of insulin-dependent anabolic signaling described above.

## Discussion

Maintenance of nutrient homeostasis depends on coordinated regulation of nutrient sensing, intracellular signaling, and metabolic gene expression. Although the insulin/TOR pathway is the principal regulator of carbohydrate and lipid metabolism in *Drosophila* (Baker & Thummel, 2007; Chatterjee & Perrimon, 2021; Teleman et al., 2005b), the upstream mechanisms controlling this pathway remain incompletely understood. In this study, we identify *Neprilysin-like 15 (Nepl15)* as a previously unrecognized regulator of nutrient partitioning that modulates insulin-dependent anabolic signaling without affecting food intake, insulin-like peptide expression, or insulin receptor abundance. Together with our previous findings that *Nepl15* mutants exhibit altered nutrient storage despite normal feeding behavior and unchanged adipokinetic hormone signaling ((Banerjee et al., 2021); Arzoo et al., 2026), these results indicate that *Nepl15* acts downstream of systemic insulin production.

A major finding of this study is that transcript levels of *Dilp2*, *Dilp5*, *Dilp6*, and *InR* remained unchanged, whereas Akt phosphorylation was markedly reduced and dFoxo abundance increased. Since Akt is the principal downstream effector of insulin signaling, reduced Akt activation would be expected to decrease mTOR activity while relieving inhibition of dFoxo, thereby suppressing anabolic metabolism (Baker & Thummel, 2007; Teleman et al., 2005a). Consistent with this model, our previous work demonstrated reduced *mTOR* expression together with increased *Thor* expression in *Nepl15* mutants (Arzoo et. al, 2026(Banerjee et al., 2021)). The additional reduction in *Glut1* expression observed in male mutants further suggests impaired cellular glucose uptake, providing a mechanistic explanation for the reduced glycogen storage previously reported in these flies.

The transcriptional changes observed in metabolic genes further support attenuation of anabolic metabolism. Male mutants exhibited reduced expression of both *GlyS* and *GlyP*, together with decreased expression of the lipogenic genes *Lipin*, *Acc*, *Fasn1*, and *Fasn2*, whereas *Midway* and *Brummer* remained unchanged. These findings suggest that *Nepl15* primarily influences glycogen and de novo lipid synthesis rather than triglyceride mobilization. Similar metabolic genes have been shown to play central roles in regulating glycogen and lipid homeostasis in *Drosophila* (Beller et al., 2008; Kühnlein, 2012; Martínez et al., 2020; Yamada, Habara, Kubo, & Nishimura, 2018). In contrast, females exhibited relatively modest transcriptional changes, including increased *Acc* expression despite reduced *Fasn1*, suggesting the presence of compensatory mechanisms that partially preserve metabolic homeostasis.

The present study also links *Nepl15* to cellular energy metabolism. We previously reported reduced *AMPK*_α_ expression in both sexes (Arzoo et al., 2026), while the present work demonstrates reduced expression of *spargel*, the *Drosophila* homolog of mammalian **PGC-1**_α_, specifically in males. Because AMPK and PGC-1α coordinately regulate mitochondrial function and metabolic adaptation (Hardie, 2011), these observations suggest that *Nepl15* influences not only insulin-dependent nutrient storage but also broader transcriptional programs governing cellular energy metabolism.

The pronounced male-specific molecular phenotype likely reflects fundamental sexual differences in nutrient allocation. Female flies continuously partition glycogen and lipid reserves toward oogenesis and embryogenesis (Sieber, Thomsen, & Spradling, 2016; Song et al., 2019), potentially buffering the effects of reduced insulin signaling. In contrast, males exhibited coordinated suppression of glycogen and lipogenic gene expression, consistent with their previously reported reductions in whole-body glycogen and triglyceride stores (Banerjee et al., 2021)

Finally, our findings extend the known metabolic functions of the neprilysin family. Mammalian neprilysin regulates numerous peptide hormones involved in glucose homeostasis, insulin sensitivity, and obesity, and elevated NEP activity has been associated with insulin resistance and metabolic disease {Turner, 2000, The neprilysin family in health and disease;Nalivaeva, 2013, Neprilysin (Chapter 127), Handbook of Proteolytic Enzymes;Standeven, 2011, Neprilysin, obesity and the metabolic syndrome;Willard, 2017, Improved glycaemia in high-fat-fed neprilysin-deficient mice is associated with reduced DPP-4 activity and increased active GLP-1 levels}. We demonstrate that *Nepl15*, a *Drosophila*-specific neprilysin, similarly influences metabolic homeostasis by modulating the insulin–Akt–Foxo–mTOR signaling axis. Collectively, these findings establish *Nepl15* as an upstream regulator of nutrient partitioning and provide new insight into how neprilysin family members coordinate intracellular signaling with glycogen and lipid metabolism.

## Au*thor* contribution

S.H.A.: Experimental design, methodology, investigation, validation, formal analysis, data curation, visualization, writing, original draft preparation, review and editing.

S.J.B.: Conceptualization, experimental design, supervision, project administration, resources, methodology, investigation, validation, formal analysis, data curation, visualization, writing, original draft preparation, review and editing, and funding acquisition.

All authors have read and agreed to the published version of the manuscript.

## Acknowledgement

All fly husbandry and molecular experiments were carried out in the Department of Biological Sciences, Texas Tech University.

